# The *SSU1* checkup, a rapid tool for detecting chromosomal rearrangements of the *Saccharomyces cerevisiae* chromosome XVI. An ecological and technological study on wine yeast

**DOI:** 10.1101/2020.04.23.055350

**Authors:** Philippe Marullo, Olivier Claisse, Maria Laura Raymond Eder, Marine Börlin, Nadine Feghali, Margaux Bernard, Jean-Luc Legras, Warren Albertin, Alberto Luis Rosa, Isabelle Masneuf-Pomarede

## Abstract

Chromosomal rearrangements (CR) such as translocations, duplications and inversions play a decisive role in the adaptation of microorganisms to specific environments. In enological *Saccharomyces cerevisiae* strains, CR involving the promoter region of the gene *SSU1* lead to a higher sulfite tolerance by enhancing the SO_2_ efflux. To date, three different *SSU1* associated CR events have been described, including translocations XV-t-XVI and VIII-t-XVI and inversion inv-XVI. In the present study, we developed a multiplex PCR method (*SSU1* check-up) that allows a rapid characterization of these three chromosomal configurations in a single experiment. Nearly 600 *S. cerevisiae* strains collected from fermented grape juice were genotyped by microsatellite markers. We demonstrated that alleles of the *SSU1* promoter are differently distributed according to the wine environment (cellar versus vineyard) and the nature of the grape juice. Moreover, rearranged *SSU1* promoters are significantly enriched among commercial starters. In addition, nearly isogenic strains collected in similar environments show different CR suggesting that translocation events occur with a non-negligible frequency in clonal populations likely due to mitotic recombination events. Finally, the link between the nature of *SSU1* promoter and the tolerance to sulfite was statistically validated in natural grape juice containing various SO_2_ concentrations. The *SSU1* check-up is therefore a convenient new tool for addressing population genetics questions and for selecting yeast strains by using molecular markers.

## 2) Introduction

Microorganisms develop various strategies for being better adapted to various environments. Among them, the yeast *Saccharomyces cerevisiae* is a noteworthy example of a microorganism whose evolution led to specialized genetic groups associated with different human-related environments (Sicard and Legras, 2011; Borneman and Pretorius, 2014; Marsit and Dequin, 2015). In a winemaking context, this species has been exposed to stressful conditions (high alcohol content, high osmotic pressure, low pH, etc.) for millennia, potentially resulting in adaptive differentiation.

In wine production, sulfite addition is widely used since the middle age as a preservative because of its antimicrobial, antioxidant, and antioxydasic activities. Produced by dissolution of sulfur dioxide (SO_2_), sulfite inhibits key glycolytic enzymes like *Tdh* and *Adh* proteins, binds carbonyl compounds such as pyruvate and acetaldehyde (Hinze and Holzer, 1986), affects transporter activity by binding membrane proteins (Divol et al., 2012), and down-regulates the expression of many central metabolism genes (Park and Hwang, 2008). Therefore, sulfite tolerance has been unconsciously selected by wine making practices and constitutes a desired trait in *Saccharomyces* wine yeast strains. Cellular mechanisms of sulfite tolerance have been extensively reviewed in *S. cerevisiae* (Divol et al., 2012; García-Ríos and Guillamón, 2019). They include the overproduction of acetaldehyde (Cheraiti et al., 2010), the regulation of sulfite reduction systems and more generally of the sulfur metabolic pathway (Divol et al., 2012). Moreover, sulfite tolerance mostly depends on the pumping of SO_2_ through the plasma membrane. This sulfite efflux involves the sulfite pump SSU1p which is encoded by the *SSU1* gene. This gene shows a high level of polymorphism (Aa et al., 2006) and deleterious mutations in its coding sequence cause SO_2_ susceptibility (Avram and Bakalinsky, 1997; Park and Bakalinsky, 2000).

The expression level of *SSU1* has a direct consequence on sulfite tolerance and has been widely studied (Pérez-Ortín et al., 2002; Nardi et al., 2010; Engle and Fay, 2012; Zimmer et al., 2014; García-Ríos et al., 2019). Interestingly, the *SSU1* promoter sequence is involved in three Chromosomal Rearrangements (CR) (i.e., XV-t-XVI, VIII-t-XVI and inv-XVI) that increase its expression leading to a more efficient sulfite pumping over (Pérez-Ortín *et al.*, 2002; Zimmer *et al.*, 2014; García-Ríos *et al.*, 2019). These three independent CR events constitute a hallmark on parallel evolutionary routes driven by human selection. In the VIII-t-XVI translocation, the native promoter of *SSU1* is replaced by tandem repeated sequences of the *ECM34* promoter from chromosome VIII (Pérez-Ortín et al., 2002). In the XV-t-XVI translocation, the upstream region of *SSU1* is placed head to tail with the *ADH1* promoter from chromosome XV (Zimmer et al., 2014). The inversion of chromosome XVI (inv-XVI) involves the *SSU1* and *GCR1* regulatory regions, increasing the expression of *SSU1* (García-Ríos *et al.*, 2019).

To date, the distribution of translocation (XV-t-XVI and VIII-t-XVI) and inversion (inv-XVI) events of the *SSU1* gene have been investigated for a small number of strains (Pérez-Ortín *et al.*, 2002; Zimmer *et al.*, 2014; García-Ríos *et al.*, 2019). Here, we set up a multiplex method (*SSU1 checkup*) based on labeled primers with different fluorochromes, to identify in a single assay the three types of *SSU1* associated CR (VIII-t-XVI, XV-t-XVI and inv-XVI) as well as the wild type forms of these chromosomes (VIII-wt, XV-wt and XVI-wt). The *SSU1 checkup* was applied to nearly 600 yeast strains, including natural isolates and commercial starters, and provides new insights on the allele frequency of rearranged *SSU1* promoters. In addition, by using microsatellite genotyping, the genetic relationships between strains of the collection were established allowing the study of CR occurrence in nearly isogenic clones. Finally, for a subset of strains, the phenotypic impact of different CR was evaluated by measuring their parameters of growth in grape juice containing different concentrations of SO_2_.

## 1) Materials and Methods

### Origin of samples

A total of 628 *S. cerevisiae* isolates were collected from grapes and fermented must (white, red and sweet) originating from five different countries (France, Lebanon, Argentina, Spain and Italy) and two different *Vitis* species (mostly *vinifera* and to a lesser extent *labrusca*) corresponding to nine different varieties. Two different procedures were used for strain isolation depending on the environment considered: vineyard or cellar. For vineyard isolates, around 2 kg of healthy and mostly undamaged grapes were collected a few days before the harvest in the vineyard, crushed in sterile conditions and macerated for 2 hours with 50 mg/L of SO_2_. The juice was fermented at 21 °C in small glass-reactors (500 mL). For cellar isolates, yeast colonies were obtained from spontaneous fermentation vats containing sulfited grape juices according to local enological practices (ranging from 20 to 50 mg/L of sulfur dioxide) except for sweet wines for which no sulfur dioxide was added. For both sampling procedures, fermentations were allowed to proceed until 2/3 of the must sugars were consumed and fermented juices were plated onto YPD plates (yeast extract, 1 % w/v; peptone, 1 % w/v; glucose, 2 % w/v; agar 2 % w/v) with 100 μg/mL of chloramphenicol and 150 μg/mL of biphenyl to delay bacterial and mold growth. Around 30 colonies per sample were randomly chosen and after sub-cloning on YPD plates, each yeast colony was stored in 30 % (v/v) glycerol at −80 °C. Additionally, a collection of 103 industrial *S. cerevisiae* starters was constituted by streaking on YPD plates a small aliquot of Active Dry Yeast obtained from different commercial suppliers.

### Microsatellite analysis

Strains were genotyped using fifteen polymorphic microsatellite loci (C3, C4, C5, C6, C8, C9, C11, SCAAT1, SCAAT2, SCAAT3, SCAAT5, SCAAT6, SCYOR267C, YKL172, YPL009) developed for estimating the genetic relationships among *S. cerevisiae* strains (Legras et al., 2007). Most of the strains were previously genotyped in our lab (Börlin *et al.*, 2016; Raymond Eder, Conti and Rosa, 2018; Emilien Peltier *et al.*, 2018;(Börlin, 2015) and the additional 82 strains were genotyped in this work using identical experimental conditions. Briefly, two multiplex PCRs were carried out in a final volume of 12.5 μL containing 6.25 μL of the Qiagen Multiplex PCR master mix (Qiagen, France), 1 μL of DNA template, and 1.94 μL of each mix, using the conditions previously reported (Peltier et al., 2018b). Both reactions were run using an initial denaturation step at 95 °C for 5 min, followed by 35 cycles of 95 °C for 30 s, 57 °C for 2min, 72 °C for 1 min, and a final extension step at 60 °C for 30 min. The size of PCR products was determined by the MWG company (Ebersberg, Germany), using 0.2 μL of 600 LIZ GeneScan (Applied Biosystems, France) as a standard marker, and chromatograms were analyzed with the GeneMarker (V2.4.0, Demo) program. Only strains that amplified at least 12 of 15 loci were kept. On the 735 strains collected, 586 met this criterion and were used in this study (listed on Table S1). The microsatellite data set was analyzed by means of the *poppr* R package using the Bruvo’s distance matrix. Strains showing a strong similarity were identified by applying a cut of value of 0.15 to the Bruvo’s genetic distance matrix. In this way, 194 very closely related strains were identified and considered as ‘clones’. This cut off value was defined in order to restore the normality of the distribution (Figure S1). The assignment of clustering methods was achieved by using the *find.clusters* function (*adegenet* package). The selection of the optimal groups was computed by the Ward’s clustering method using the Bayesian Information Criterion (BIC) as statistical criterion (Avramova et al., 2018).

### The SSU1-checkup method

In order to experimentally detect in a single multiplex PCR test all the CR involving the gene *SSU1*, labeled primers were designed using a specific dye per chromosome position as follows: 6-FAM (Chr8: VIII-14558), ATTO550 (Chr15: XV-160994), HEX (Chr16: XVI-373707) and ATTO565 (Chr16: XVI-412453) (Table 1). All the primers (Tables 1 and S2) were synthesized by Eurofins genomics (Ebersberg, Germany). A multiplex PCR was carried out in a final volume of 20 μL using 100 nM of each primer, 1 μL of template DNA and the Qiagen PCR multiplex PCR kit (Qiagen, France) on a T100TM Thermal cycler (Bio-Rad, France). The following PCR program allows the amplification of all the expected fragments from the rearranged and the wild type VIII, XV and XVI chromosomes: initial denaturation at 95 °C for 15 min, followed by 35 cycles of 94 °C for 30 s, 55 °C for 90 s, 72 °C for 90 s, ending with a hold at 60 °C for 30 min. DNA templates for PCR were extracted in 96-well microplates using the previously described LiAc-SDS protocol (Chernova et al., 2018). Before analysis, PCR products were diluted 60 times in ddH_2_O and 1 μL of this solution was mixed with 0.2 μL of the internal size standard GenScanTM 1200 LIZ^®^ (Applied Biosystems, France) and 9.8 μL of highly deionized Hi-DiTM formamide (Applied Biosystems, France). Samples were analyzed by Eurofins genomics (Ebersberg, Germany) on an ABI-3710 Genetic Analyzer. Each peak was identified according to the color and size and attributed to the alleles (Figure 1). Each allele was also sequenced by amplifying both strands with non-labeled primers. The sequences were released on GenBank with the following accession numbers: ID MT028493-MT028507.

**Table 1.**
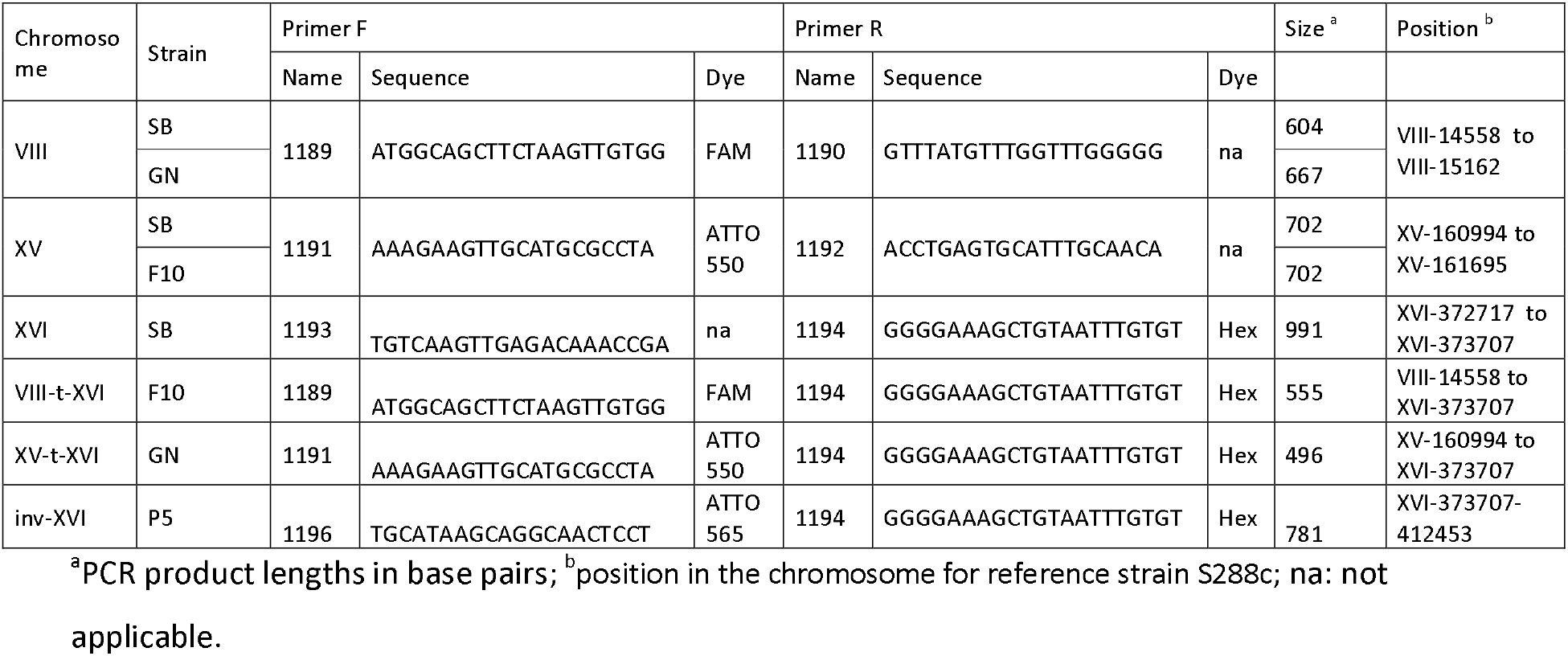
Primers used for the SSU1 checkup method and the strains used as positive controls

**Figure 1.**
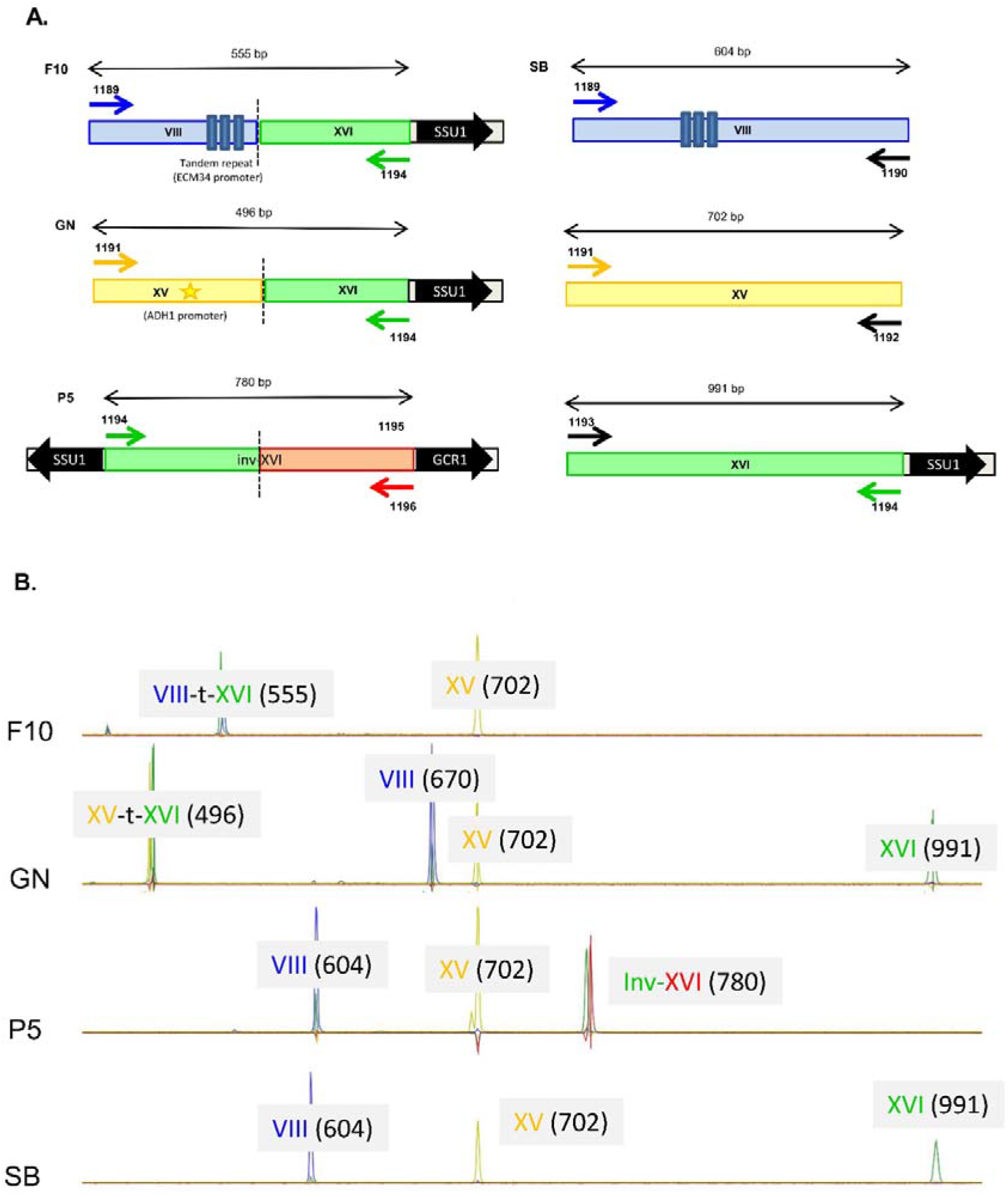
Panel A. Schematic positioning of primers used for SSU1 checkup, blue, yellow, green, and red colors represent the specific dye used for each chromosome, FAM, ATO550, HEX and ATO 565. The relative position of the gene SSU1 and the modified promoter regions (ECM34, ADH1 and GCR1) are indicated. The hatched line represents the chromosomal break point in rearranged strains GN, F10 and P5. The panel B represents chromatograms of multiplexed PCR reactions for the reference strains. The wt and rearranged XVI chromosomes were detected by co migration of fragment of different size labeled with specific dyes. For Chromosome VIII different sizes were obtained according to the strain due to the differential number of tandem repeats in ECM34 promoter.

### SO_2_ tolerance assessment

To assess SO_2_ tolerance, a subset of 34 strains (Table S3) were cultivated in white grape juice (Sauvignon Blanc from Bordeaux area, France). This must had a total SO_2_ concentration of 14mg/L and was spiked with 0, 25, 50, and 75 mg/L of total SO_2_. Cultures were achieved in 96-well plates (U flat well, Greiner, France) filled with 200 μL of grape juice sterilized by a nitrate-cellulose membrane filtration (Millipore, France). Yeasts were pre-cultivated in YPD media (yeast extract, 1 % w/v; peptone, 1 % w/v; glucose, 2 % w/v) for 16 h at 28 °C and inoculated into the grape juice to a final concentration of 1×10^6^ cells/mL. Growth was monitored by OD_600_ measurements for 96 hours at 28°C using a microplate spectrophotometer (Synergy HT Multi-Mode Reader, BioTek Instruments, Inc, USA). Culture plates were shaken every 25 minutes for 30 s prior to the OD_600_ measurements. The well position on the microplate was randomized and six replicates were done for each strain*media condition. Data from the microplate reader were transformed with the polynomial curve y=−0.0018*×3+0.1464*×2+0.7757*×+0.0386 to correct the non-linearity of the optical recording at higher cell densities as previously reported (Martí-Raga et al., 2016). Growth kinetic data were fitted using the *Richards* flexible inflection point model implemented by the *fit growthmodel* function, R package *growthrates*. This model allows the estimation of the maximal growth rate (*μmax*). A second parameter, *Lag Time*, was manually computed from raw data by considering the time necessary to reach twice the OD_600_ of the inoculum. A linear model was applied for estimating effects of the SO_2_ concentration and type of chromosome XVI and their possible interactions:

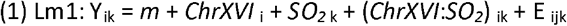

Where Y are the values of the trait (*μmax* and *Lag Time*), for *j ChrXVI* configurations _(i=1 to 6)_, and *k SO_2(k=1 to 4)_* concentrations, *m* was the overall mean and *E*_*ijk*_ the residual error. Homoscedasticity of the ANOVA was tested by *LeveneTest* function (*car* package) while the normal distribution of models’ residuals was estimated by visual inspection (*qq plot*).

## 4) Results and discussion

### Assessment of the genetic diversity of starters and natural isolates populations of *S. cerevisiae*

In this study we analyzed a large dataset of 586 isolates that were genotyped using 15 microsatellite loci. This collection includes 103 industrial starters and 483 indigenous isolates from different origins (sampling mode, red or white grape must, country). Since many natural isolates were sampled in the same juice, some of them could have originated from clonal expansion and be very similar from a genetic point of view. A filtering procedure was applied for keeping only one representative genotype of each clonal population by removing all but one strain having a Bruvo’s genetic distance lower than 0.15 (Figure S1). By this procedure, many isolates closely related to industrial starters were identified (Figure S2) demonstrating the wide dissemination of commercial yeasts in vineyard and winery environments as previously reported (Valero et al., 2005; Börlin, 2015). In addition, 21 isogenic strains were found among commercial starters. This filtering procedure defined three subpopulations: “starters = 82”, “natural isolates = 310” and “closely related clones = 194” (Table S1). The genetic relationships for each strain within the *starters* and *natural isolates* subpopulations were then analyzed by a principal component analysis (k=6). The constitution of genetic groups based on microsatellites inheritance was carried out by using a k-mean based algorithm (see Methods).

This genetic analysis clustered the 82 commercial strains in three groups with a group C clearly separated from the other two (Figure S3) and corresponding to “Champenoise” strains, a particular wine yeast group previously described (Legras et al., 2007; Novo et al., 2009; Borneman et al., 2016). Its detection validated our clustering analysis based on the use of k-mean clustering. The structure of the 310 *natural isolates* collected was also investigated and six subgroups were defined. Figure 2A shows the first two dimensions of the PCA; axis one clearly identified a group of isolates from Argentina, while axis 2 broadly discriminated the five other groups. The assignment of subgroups on neighbor-joining tree (unrooted) illustrates that isolates are mostly clustered according to their geographical origins (Figure 2B). Some groups are specific to sampling zones such as group 3 (n=16) and group 6 (n=84) that only contain strains sampled in Argentina and Lebanon, respectively. In contrast, group 2 (n=177) encompassed isolates from different geographic origins (Italy, Spain, France and Lebanon) (Table S4). Although not perfectly discriminating, this first analysis filtered the redundancy of our collection and provided a clear overview of the genetic diversity of non-redundant strains.

**Figure 2.**
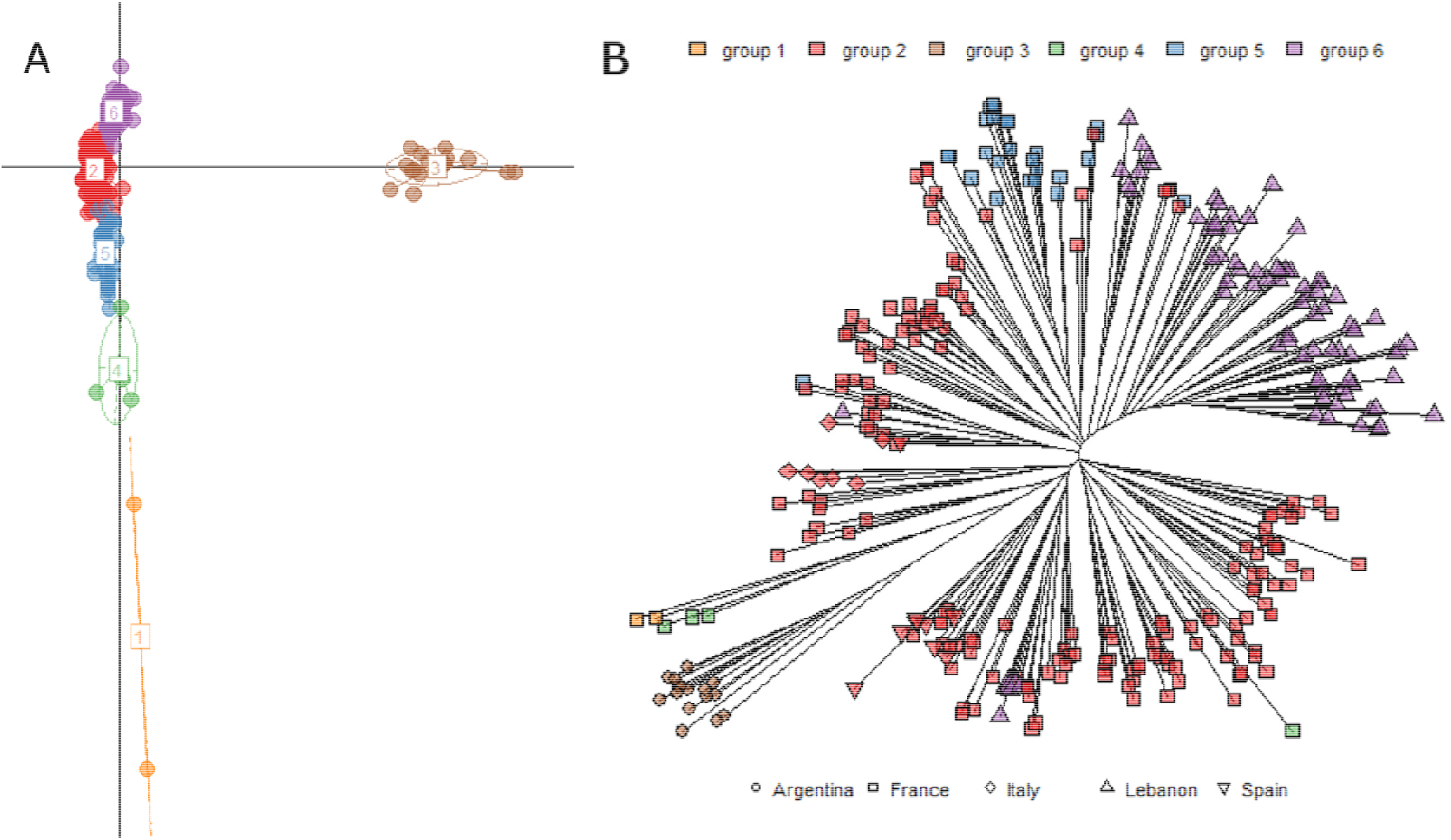
Panel A. Principal component analysis of 310 natural isolates discriminated by 15 polymorphic loci. The seven groups represented (1 to 6) were inferred by k-mean clustering. The panel B represents the position of the strains according to the inferred groups and the country origin of their sampling.

### Development of the *SSU1* checkup, a simple method for genotyping *SSU1* chromosomal rearrangements in *S. cerevisiae*

To date, translocation events (i.e., XV-t-XVI and VIII-t-XVI) have been detected by classical PCR experiments narrowing the chromosomal break points identified by two original works (Pérez-Ortín et al., 2002; Zimmer et al., 2014). Recently, an additional chromosomal rearrangement involving the gene *SSU1* (inv-XVI) was also described (García-Ríos et al., 2019). Since these classical PCR amplifications are scarcely adapted to screen multiple genotypes in large populations, a multiplexed method (*SSU1 checkup*) was set up aiming to identify, in a single PCR reaction, these three types of chromosomal rearrangements (VIII-t-XVI, XV-t-XVI and inv-XVI) as well as the wild type alleles of the corresponding chromosomes (VIII-wt, XV-wt and XVI-wt). Labeled primers harboring different fluorochromes allowed the rapid identification of allele sizes ranging between 388 and 991 bp. In preliminary studies, we used reference strains SB (XVI-wt), GN (XV-t-XVI), F10 (VIII-t-XVI), and P5 (inv-XVI), to design and validate primers. Primers position, as well as the length of the DNA fragments amplified for the reference strains, are summarized in Table 1. The sequences of all the alleles identified were submitted to GenBank (ID 2310529). Figure 1 presents the schematic position of each primer and the chromatograms obtained for each reference strain. In order to simplify the chromatographic patterns, the detection of reciprocal translocation events (XVI-t-XV and XVI-t-VIII) was not included in the *SSU1 checkup*. Primers used to amplify these reciprocal translocations are given in Table S2.

### Landscape of the different *SSU1*-promoter alleles in natural and selected populations

Initially, we developed the *SSU1* checkup to track two translocation events (VIII-t-XVI and XV-t-XVI). In the VIII-t-XVI translocation, the native promoter of *SSU1* is replaced by DNA sequences of the *ECM34* promoter (located on chromosome VIII) (Pérez-Ortín et al., 2002). Alleles for this CR (i.e., VIII-t-XVI^388^, VIII-t-XVI^478^, VIII-t-XVI^555^ and VIII-t-XVI^631^ bp) result from the alternative number of units of tandem repeated motifs (76 bp and/or 47 bp) localized in the promoter region of the gene *ECM34* (Figure S4). In the XV-t-XVI translocation, the upstream region of *SSU1* is placed head to tail with the *ADH1* promoter (located on chromosome XV) (Zimmer et al., 2014) and a single allele has been recognized (XV-t-XV^496^) (Figure 1B). Surprisingly, for some natural isolates, we did not detect either the XVI-wt allele or the translocated alleles. These unexpected outcomes could have been the result of a potential PCR artifact (i.e., limited primer annealing or a possible synteny modification linked to the *SSU1* gene). However, 39 strains failed to amplify the expected DNA fragments, even when performing single PCR reactions using alternative primers (Table S2). These experiments discarded the hypothesis of a partial annealing of primers. The recent report of a new chromosomal rearrangement (inv-XVI) involving the gene *SSU1* (García-Ríos et al., 2019), prompted us to include this new allelic form of the gene *SSU1* to our screening (see Figure 1). Of the 39 strains concerned, this chromosomal inversion was detected in 19 natural isolates and showed a single allele of 781 bp (inv-XVI^781^). The remaining 20 strains definitively failed to amplify any fragment. This suggests that these strains could harbor another still uncharacterized chromosomal rearrangement flanking the *SSU1* gene. The implementation of the additional inv-XVI-associated primer completed the *SSU1-checkup* method, which can detect four types of chromosome XVI structures in a single PCR reaction.

Since the amplified fragments are physically linked to the chromosome XVI’s centromere (*CEN16*), they belong to chromosome XVI during the cell division process. Therefore, native and rearranged chromosome XVI alleles were merged in order to have an integrated overview of the chromosome XVI inheritance. This allow to count possible case of extra copy number of chromosome XVI (aneuploidy) due to CR. The allele frequencies computed are given in Table 2; the occurrence of each allele between *natural isolates* and *starters* populations was compared by a Chi^2^ test. Among the 392 non-redundant *S. cerevisiae* strains analyzed, the most frequent alleles found were VIII-t-XVI ^555^ (0.41) and XVI-wt^991^ (0.34). However, their allelic frequencies are not evenly distributed. Indeed, the *starters* group (n=82) is significantly enriched in alleles VIII-t-XVI^388^ and XV-t-XVI^496^ respect to the *natural isolates* group (n=310); in contrast, *natural isolates* mostly harbor the VIII-t-XVI^555^ allele.

**Table 2.** Allele frequency and percentage of homozygosity of different chromosome XVI forms within starters and natural isolates populations

| Chromosome XVI alleles | Allele frequencies |  |  |  | % of homozygous strains |  |  |  |
| --- | --- | --- | --- | --- | --- | --- | --- | --- |
|  | Total samples n=392 | Natural isolates n=310 | Starters n=82 | Chi<sup>2</sup> test | Total samples n=392 | Natural isolates n=310 | Starters n=82 | Chi<sup>2</sup> test |
| VIII-t-XVI<sup>388</sup> | 0.093 | 0.021 | 0.369 | *** | 3.8 | 2.1 | 10.0 | *** |
| VIII-t-XVI<sup>478</sup> | 0.027 | 0.034 | 0.000 | nr | 2.2 | 2.8 | 0.0 | nr |
| VIII-t-XVI<sup>555</sup> | 0.411 | 0.464 | 0.206 | *** | 34.6 | 43.2 | 3.8 | *** |
| VIII-t-XVI<sup>631</sup> | 0.008 | 0.010 | 0.000 | nr | 0.8 | 1.0 | 0.0 | nr |
| VIII-t-XVI<sup>(all alleles)</sup> | 0.539 | 0.529 | 0.575 | ns | 41.4 | 49.1 | 13.8 | *** |
| XV-t-XVI<sup>496</sup> | 0.041 | 0.026 | 0.100 | *** | 1.4 | 0.7 | 3.8 | *** |
| inv-XVI<sup>781</sup> | 0.042 | 0.053 | 0.000 | nr | 4.4 | 5.6 | 0.0 | nr |
| XVI-wt<sup>991</sup> | 0.337 | 0.322 | 0.394 | ns | 21.8 | 25.4 | 8.8 | ** |

\*, \*\* and \*\*\* symbols stand for pvalue< 0.05, 0.01 and 0.005, respectively
nr: not relevant, ns: not significative

Since chromosomal rearrangements lead to more active *SSU1* genes, these alleles are supposed to be mostly dominant (Clowers et al., 2015; Peltier et al., 2018a). Therefore, for having a more accurate understanding of the functional impact of chromosome XVI forms, the percentage of homozygous strains for this locus is also given in Table 2. The homozygosity level of VIII-t-XVI alleles is much higher among *natural isolates* (49.1 % vs. 13.8 %) than among *starters*. Intriguingly, this significant discrepancy was not found for the overall homozygosity level measured for the 15 microsatellite markers with 0.62 vs. 0.57 for the *starter* and *natural isolates* groups, respectively. In the same way, although allele frequencies of XVI-wt^991^ are quite similar between the two populations, homozygous strains are less frequent in the *starters* group (8.8 % vs. 25.4 %, corrected Chi^2^ test, p=1.10^−4^). Consequently, only seven industrial strains lacked any rearranged *SSU1* allele (*ECM34-SSU1*, *ADH1-SSU1*, or *GCR1-SSU1*).

Among the 586 strains typed using the *SSU1 checkup*, 18 biallelic combinations of the seven chromosome XVI alleles were found. The frequency of isolates carrying at least one type of CR is shown in Figure 3. Industrial strains are significantly enriched in translocations VIII-t-XVI and XV-t-XVI compared to natural isolates. In contrast, the inv-XVI allele was rarer and never found in industrial strains (the reference strain P5, was not included here). Interestingly, two industrial starters (3 %) carry both translocated chromosomes. In addition, 11 starters (13 %) have an extra copy of chromosome XVI (aneuploidy), a fraction much higher than for the *natural isolates* group (1 out of 310). Some starters recommended for red grape juice winemaking or Cognac distillation (where the SO_2_ pressure is lower than for white wine) are “not rearranged”. Altogether, these results are in agreement with the hypothesis that the addition of SO_2_ in grape juices by winemakers promotes the selection of a rearranged chromosome XVI form (Zimmer et al., 2014; Clowers et al., 2015).

**Figure 3.**
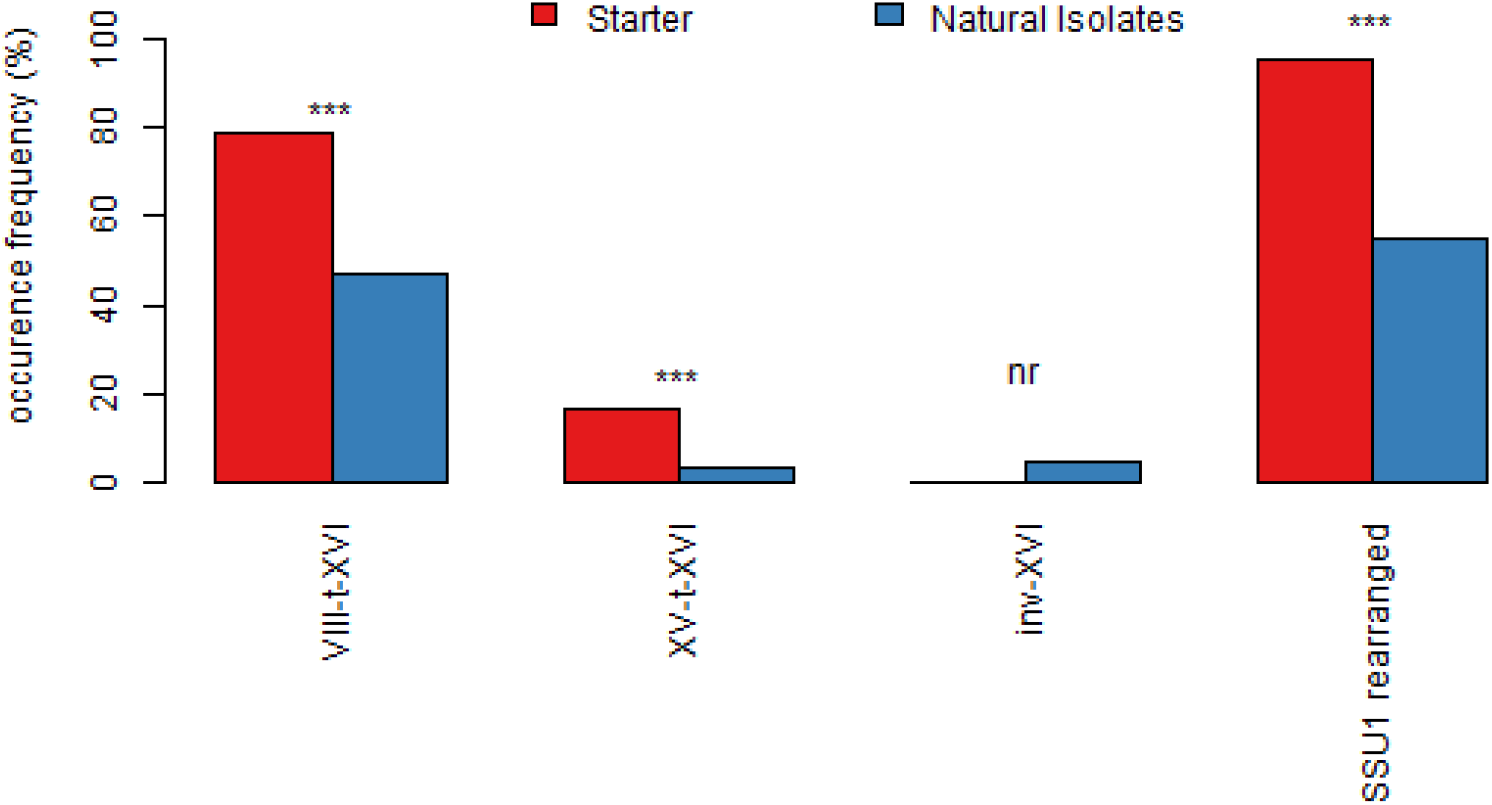
Starters are enriched in rearranged chromosome XVI forms respect to natural isolates. The frequency (%) of “Starter” (red) and “Natural” (blue) isolates carrying at least one Chromosomal Rearrangement (CR) is represented. Significant differences are marked with ***.

### Linking chromosomal rearrangement events of chromosome XVI and yeast ecology

Wine-related natural isolates allow the study of broad ecological factors influencing the chromosomal configurations of *SSU1’s* promoter. It has been shown that *SSU1* related translocations in wine yeast isolates are advantageous for growth in sulfited grape juice and contribute to a fitness gain respect to oak yeast strains (Clowers et al., 2015). Furthermore, previous studies revealed that variations in the promoter region of *SSU1* gene in wine yeasts enhance the *SSU1* gene expression during fermentation, and have a remarkable effect on the SO_2_ resistance levels (Pérez-Ortín et al., 2002; Zimmer et al., 2014; García-Ríos and Guillamón, 2019).The subpopulation of 310 unique strains characterized in this work were split according to the sampling procedure applied: *cellar* (n=205) vs *vineyard* (n=105) isolates (Table S1). *Cellar* strains were isolated from spontaneously fermented vats in various wine estates, from sulfited grape musts according to the recommended practices of the area of origin. *Vineyard* strains were isolated from grapes manually harvested, crushed and fermented in sterile laboratory conditions (see Methods).

The occurrence frequency of the 18 allelic combinations found in both groups is shown in Figure 4A. The proportion of genotypes (555:555 and, 991:555) was significantly higher in the *cellar* group while *vineyard* isolates are slightly enriched in 781:781 genotypes (p value < 0.1, corrected Chi^2^ test). Moreover, strains having inherited at least one rearranged chromosome XVI are significantly more frequent in the *cellar* group (Figure 4A). The allelic frequency observed here could reflect that among wine-related yeast isolates, *cellar* strains undergo a stronger selective pressure than *vineyard* strains likely due to winemaking operations.

**Figure 4.**
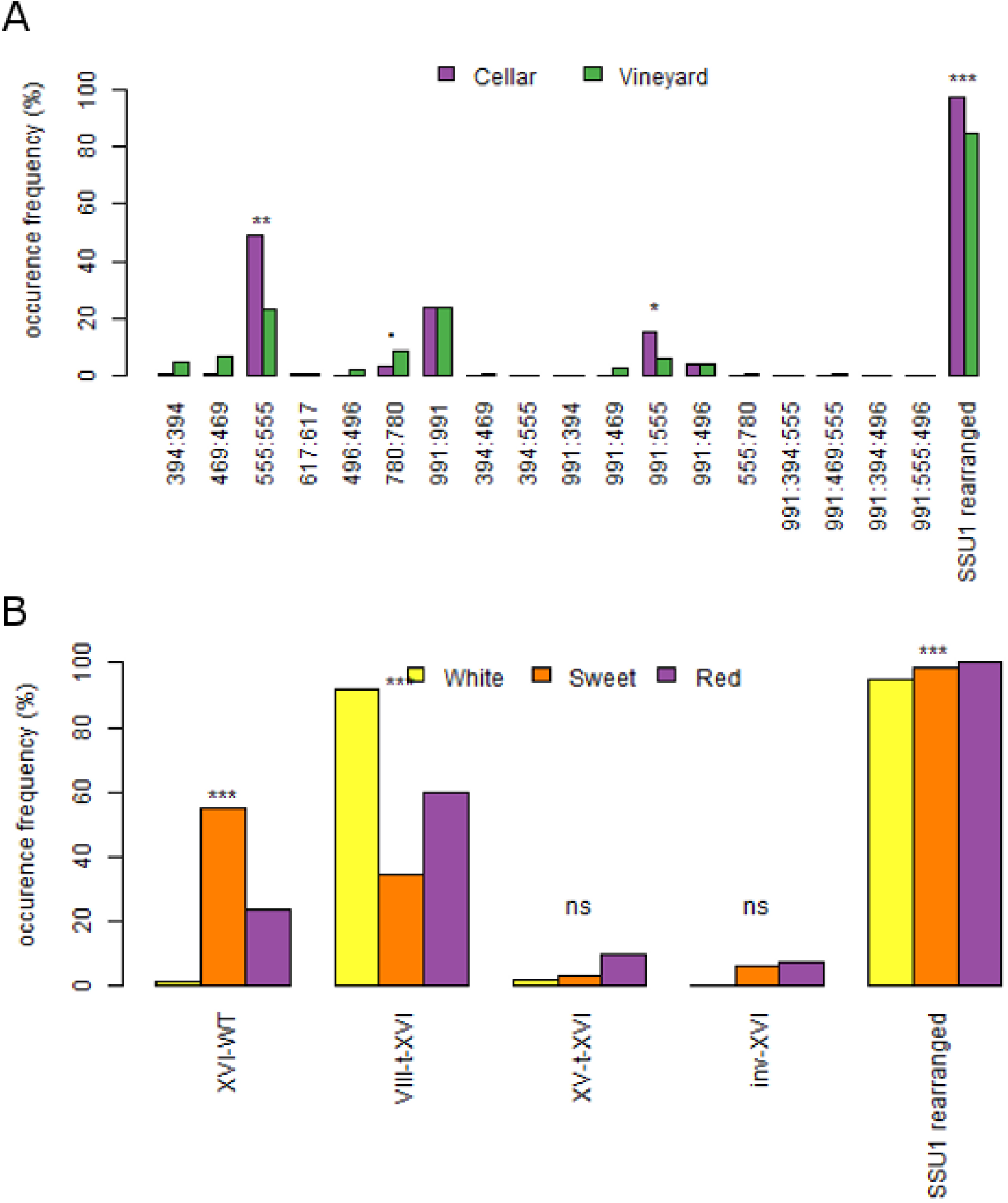
Grape juice matrixes and sampling methods impact the occurrence of SSU1 alleles. Panel A. Occurrence frequency (%) of 18 biallelic combinations of the seven chromosome XVI alleles. Panel B. Occurrence frequency (%) of rearranged chromosome XVI forms and chromosome XVI-wt form based on the type of grape juice matrix (sweet, white and red). *** significance level < 0.001; ** significance level < 0.05; * significance level < 0.1, corrected Chi^2^ test; ns: not significant.

The impact of the nature of grape juice from which strains were isolated was also tested. For this study, we focused our investigation only on *cellar* populations (n=205) that have been submitted to *in situ* enological treatments. Indeed, according to the enological practices and doses, grape juices are not equally sulfited. This is consistent with the recommendations of International Organization of the Vine and Wine (OIV) that regulates the limits for total SO_2_ in wines (150 mg/L for red wines and 200 mg/L for white wines and rosés) (OIV, 2019). Strains were split in three groups depending on the type of grape juice matrix: *sweet* (n=67), *white* (n=95) and *red* (n=43). As shown in Figure 4B, strains isolated from *white juice* are enriched in the VIII-t-XVI rearrangements and a few of them are homozygous for the XVI-wt^991^ allele. In contrast, strains isolated from *sweet grape* juices are strongly enriched in the native chromosome form. For this group, the allele frequency of XVI-wt is 0.63, which is twice as much as the frequency found for the overall population (0.33). These results are consistent with the traditional enological practices used in the Bordeaux area, where the addition of sulfite in the must is routinely used for dry white wine fermentation, but mostly avoided in the beginning of sweet wine fermentation to limit SO_2_ binding phenomena. This suggests that the selection of CR is strongly influenced by the winemaking practices used in cellars.

### Estimation of mutation frequency of the SSU1 promoter region by comparing clonal populations

The impact of translocations in the phenotypic adaptation of yeast has been widely investigated by using genetically engineered strains (Tosato and Bruschi, 2015; Fleiss et al., 2019). However, the description of spontaneous chromosomal rearrangements in clonal populations is much less described. The *SSU1*-checkup method provides an indirect opportunity to address the mutation frequency of XVI-forms in clonal populations. In this section, the pool of 194 closely related clones was used in order to identify nearly isogenic groups of strains. In order to minimize the genetic distance inside a group, only strains showing less than two VNTR (Variable Number of Tandem Repeat) or LOH (Loss of Heterozygosity) were grouped together. By this way, 16 nearly genetic groups were identified encompassing 125 strains (Table S5). Group sizes ranged between 3 and 22 individuals, with a Bruvo’s genetic distance between the strains of each group always lower than 0.106. The relative distance between each group was illustrated by a Principal Component Analysis (Figure 5A).

**Figure 5.**
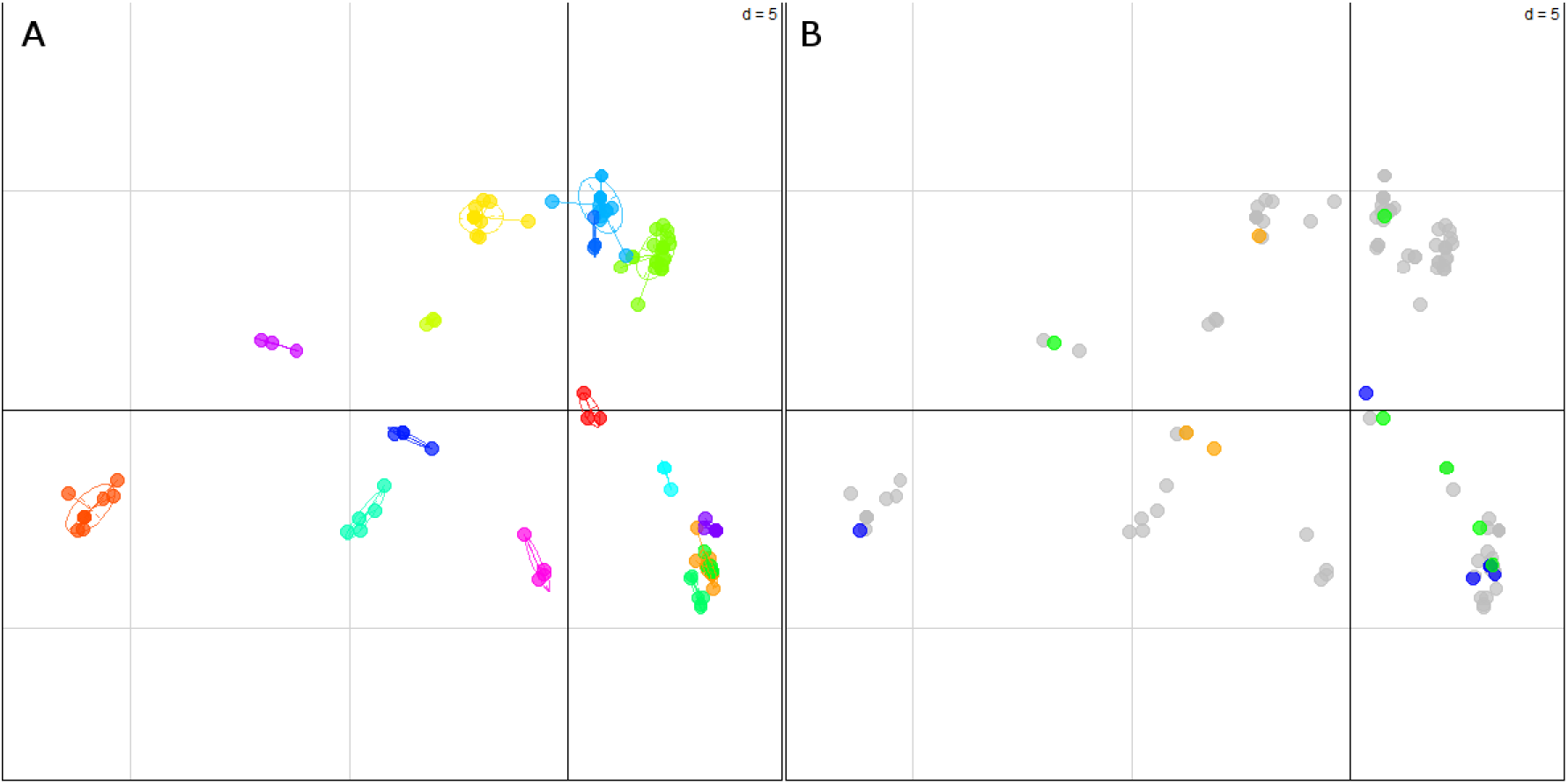
Alleles of chromosome XVI show a rapid evolution in isogenic populations. Panel A. Principal Component Analysis of 125 closely related clones (nearly isogenic) discriminated by 15 polymorphic loci. The sixteen groups were inferred manually by considering as isogenic the strains having an identical genotype for at least 13 microsatellite loci. The projection of each strain according to the 16 groups represents 14.3 % of the total inertia. The Bruvo’s genetic distance within each group is always lower than 0.105. Panel B represents the same projection, but strains were colored according to the type of change occurring on chromosome XVI. Grey dots represent the major allelic form found in each subgroup while orange, blue, and green dots represent LOH, VNTR and CR, respectively.

Each isogenic group could be therefore considered as a clonal population derived from a common ancestor with a unique genotype that has evolved in closely related individuals. According to this hypothesis, the residual genetic variability observed between individuals of the same group should result from spontaneous mutations that have occurred in the common ancestor. This residual variability was interrogated for three types of loci: microsatellites, chromosome VIII and chromosome XVI. For each group, we used as a reference the most frequent genotype (Table S5). For microsatellite loci, few VNTR and LOH mutations were detected within individuals. This is consistent with the fact that strains were mostly isolated from the same vat/cellar/area samples. Within the 15 microsatellite markers, the average mutation frequency was 2.4 % and 2.1 %, for VNTR and LOH, respectively (Table 3). In the same way, the mutation frequencies for chromosomes VIII and XVI were computed. Interestingly, a noteworthy variability was found for 10 out the 16 groups. For chromosome VIII, LOH events were observed only in groups 4 and 14 while tandem repeat shifts of *ECM34* promoter (VNTR) impacted in 4 groups. The overall allelic variability for the chromosome VIII locus is 4.5%, which is slightly higher than the average variability observed for neutral markers (microsatellites).

**Table 3.**
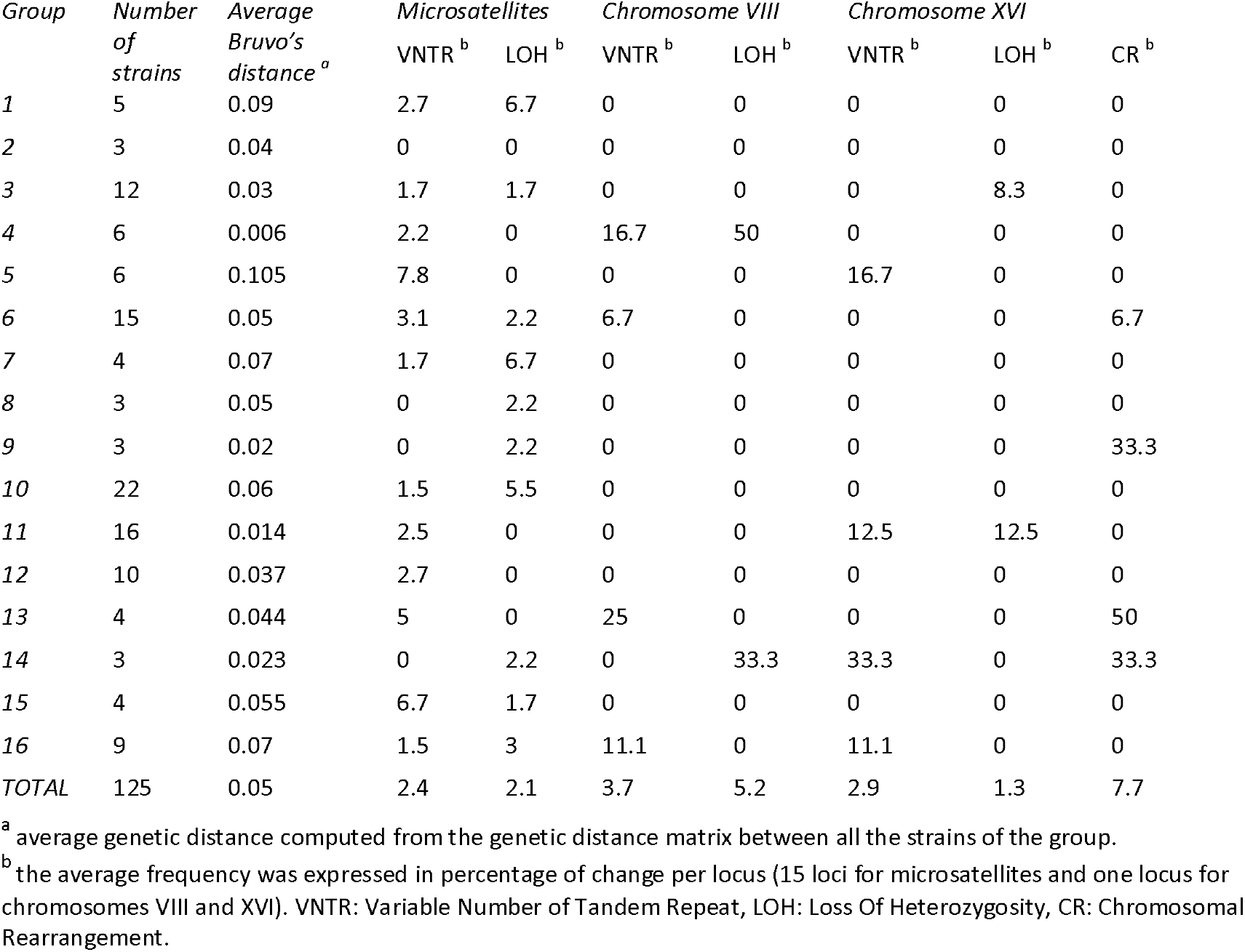
Frequency of VNTR, LOH and Chromosomal rearrangements in isogenic populations

The inheritance of chromosome XVI is more complex to analyze due to the combined influences of VNTR, LOH and the different CR (Figure 5B). These changes were observed in 7 out the 16 groups, being their overall frequency 2.9 %, 1.3 % and 7.7 %, respectively. These allelic modifications were mostly found within strains isolated at the same place and showing the same microsatellite pattern. For VNTR, four isogenic groups showed allelic variations in the VIII-t-XVI translocation. These variations are due to tandem repeats on *ECM34* promoter; their frequencies are similar to those observed for chromosome VIII (2.9% vs 3.7%). LOH variations were observed for two groups (i.e. 3 and 11). For instance, the main genotype observed in the group 11 is 555:555 (12 out 16 strains) but two strains (*3bibi6_26* and *3bebi3_13*) have the genotype 991:555. Although it is not possible to determinate the genotype of their common ancestor, this difference suggested that the main group underwent a possible LOH event. Strikingly, CR were also observed resulting in a switch of translocation form. This is the case of strains 13AQGUICUV1 (555:555) and 14AQGUICUV1 (991:496) isolated in the same area in sweet wines. The first is homozygous with a VIII-t-XVI form, the second is heterozygous with both XVI-wt and XV-t-XVI forms. A similar switch was also observed for strains 2duSPO9 (555:555) and 3mabi1_10 (991:496) that belong to group 6. These switches of chromosomal rearrangements might be due to recombination events that could have occurred in the lineage of the common ancestor. Interestingly, the strain 2duSPO9 (555:555) was isolated from a Sauvignon blanc juice while all the other strains of this group were isolated from red grape juice (Table S5). This is consistent with the fact that the translocation VIII-t-XVI^555^ was more frequently found in samples isolated from white grape juice (Figure 4B). All these findings were verified by simple PCR reactions, after additional DNA extractions. Although the number of events observed is not sufficient for providing statistically robust data, our results suggest that chromosomal rearrangements can occur with a non-negligible frequency from a common ancestor. These microevolutionary changes between an industrial strain and its descendants selected after persistence in nature were previously reported using inter-*delta* markers (Franco-Duarte *et al.*, 2015). In the case of the *SSU1* promoter, these rearrangements could play a significant role in the adaptative responses to different environments especially due to the use or not of SO_2_ in the early stages of vinification.

### Impact of chromosomal rearrangements on yeast fitness parameters in sulfited grape juice

Finally, we compared groups of unrelated strains harboring six different promoter regions of the gene *SSU1*. Five representative yeast strains of each group were selected by choosing strains with contrasted microsatellite inheritance and sampling origins. Indeed, strains belonging to each group showed an average Bruvo’s genetic distance higher than 0.50. These genetic distances were similar to those observed for the total population (Figure S5). Therefore, the strains selected may be considered as genetically unrelated. Growth kinetics of these strains in filtered Sauvignon blanc grape juice spiked with different SO_2_ concentrations were analyzed by OD_600_ measurement. Since some strains did not reach an OD_600_ plateau after 96 hours, only *μmax* (maximal growth rate) and *Lag Time* (Lag phase time) parameters were analyzed. An overview of the kinetics for the reference strains GN, SB, P5 and Fx10 is given in Figure S6. The effect of SO_2_ concentration and type of chromosome XVI were estimated by a two-way ANOVA (model Lm1, see methods). The variance explained by factors is given in Table 4. As expected, SO_2_ addition to the grape juice significantly impacted the *μmax* and the *Lag Time* parameters explaining 14.8 % and 18.4 % of total variance, respectively. This confirms the selective pressure imposed by increasing SO_2_ concentrations, which delayed the beginning of exponential growth and reduced the maximum growth rates of *S. cerevisiae* strains. In addition, the type of chromosome XVI significantly impacted these two parameters (Table 4) contributing in higher proportion to the total variance observed (23.5 % for *Lag Time* and 17.0 % for *μmax*). Finally, a significant interaction was detected between SO_2_ concentration and the type of chromosome XVI type for the *Lag Time* parameter (p < 1.10^−6^).

**Table 4.** Analysis of variance of SO_2_ and SSU1 promoter forms for lag time and μmax

| Trait | SO<sub>2</sub> |  | Chr XVI |  | SO<sub>2</sub>:Chr XVI interactions |  | Residual | Levene test |
| --- | --- | --- | --- | --- | --- | --- | --- | --- |
|  | Effect | p value <sup>a</sup> | effect | p value <sup>a</sup> | effect | p value |  |  |
| Lag time | 18.4 | *** | 23.5 | *** | 7.0 | *** | 50.7 | 0.06 |
| μ<sub>max</sub> | 14.8 | *** | 17.0 | *** | 2.5 | ns | 65.6 | 0.002 |

\*\*\* significance level < 1.10<sup>-6</sup>; ns: not significant.

The specific impact of the form of the *SSU1* promoter is illustrated in Figure 6. Since five unrelated strains were tested in each group, the impact of the different *SSU1* promoters is partially decoupled to the strain effect. As expected, *“non-rearranged strains”* (i.e., XVI-wt/XVI-wt) present the highest *Lag Time* and the lowest *μmax* at high SO_2_ concentrations. The translocation XV-t-XVI and the inversion (inv-XVI) appear to be very efficient to cope with higher SO_2_ concentrations. Indeed, strains carrying these alleles were poorly affected by the addition of 75 mg/L of SO_2_. These findings are consistent with the elevated *SSU1* expression levels reported for these two chromosomal rearrangements (Zimmer et al., 2014; García-Ríos and Guillamón, 2019). For the VIII-t-XVI translocations, only three allelic forms (VIII-t-XVI^388^, VIII-t-XVI^478^, VIII-t-XVI^555^) were tested. The fourth allele (VIII-t-XVI^631^) was found in only three strains, two of them being clearly isogenic. The allele VIII-t-XVI^478^ was the most efficient in reducing the *Lag Time* and preserving the *μmax*. The other two alleles (VIII-t-XVI^388^ and VIII-t-XVI^555^) resulted in the least favorable rearrangements for sulfite tolerance (Figure 6).

**Figure 6.**
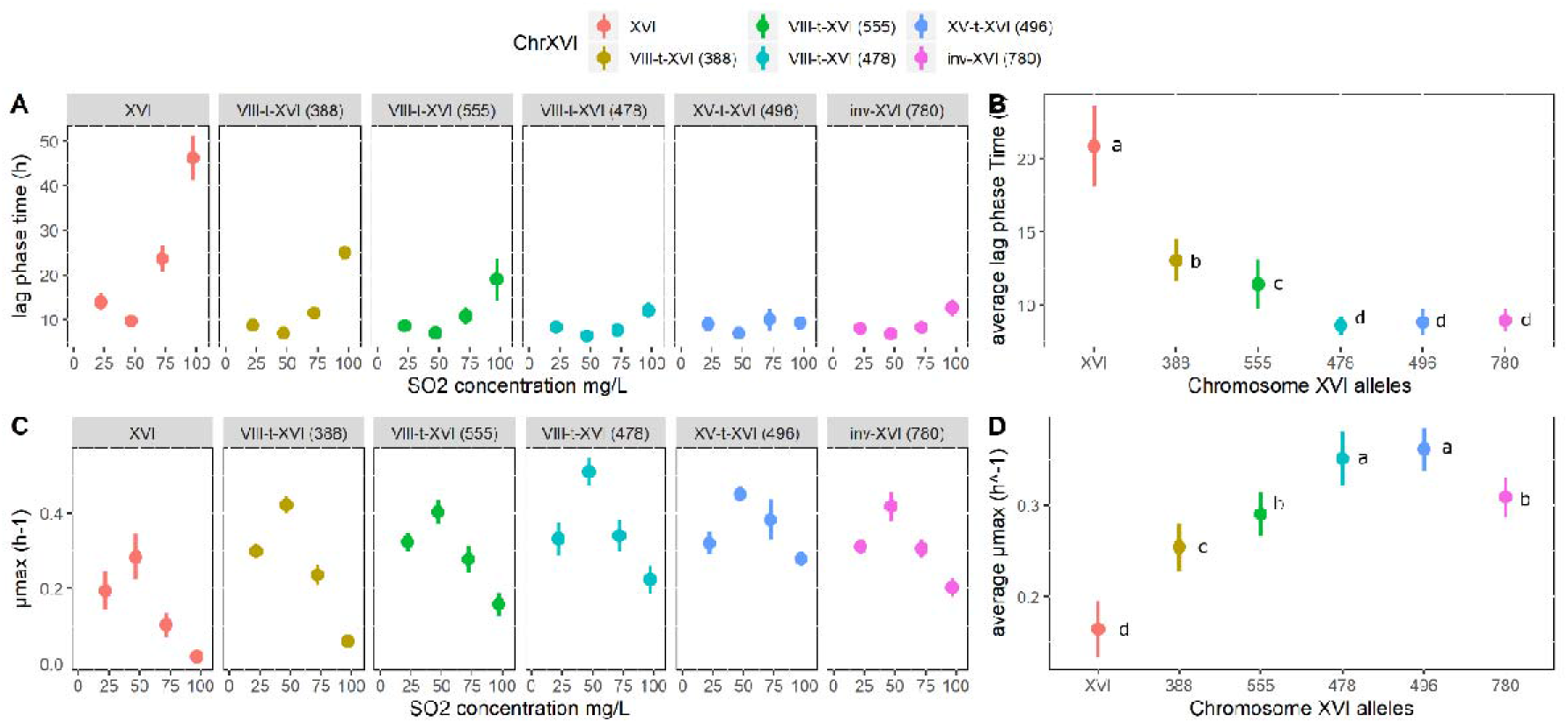
Phenotypic impact of six SSU1 promoter alleles. Panel A. Mean of the Lag Times (in hours) of the 5 strains tested at different SO_2_ concentrations, for each of the chromosome XVI alleles. Panel B. Mean of the Lag Times (in hours) of the 5 strains tested at all SO_2_ concentrations, for each of the chromosome XVI alleles. Panel C. Mean of the μmax of the 5 strains tested at different SO_2_ concentrations, for each of the chromosome XVI alleles. Panel D. Mean of the μmax of the 5 strains tested at all SO_2_ concentrations, for each of the chromosome XVI alleles.

Differences in the length of the VIII-t-XVI translocation alleles mainly result from the variable number of 76 bp tandem repeat units (Goto-Yamamoto et al., 1998). The number of tandem repeat units (76 bp and 47 bp) for the VIII-t-XVI translocation was verified by DNA sequencing and results are summarized in Figure S4. Alleles VIII-t-XVI^388^, VIII-t-XVI^478^, and VIII-t-XVI^555^ show two, three, and four 76 bp tandem repeats plus one additional 47 bp tandem repeat, respectively. Since the allele VIII-t-XVI^478^ out competed the other two forms (Figure 6), we can conclude that the number of tandem repeats and the SO_2_ tolerance are not fully correlated, as previously reported (YUASA et al., 2004). Therefore, undetermined natural allelic variations in the *SSU1* coding sequence, in flanking regions, or in other genomic loci are possibly also involved in this trait.

By comparing for the first time the effect of six configurations of the *SSU1* promoter on yeast fitness we pave the avenue for screening sulfite tolerance of strains in a simple genetic test. At first glance, the translocation XV-t-XVI and the inversion (inv-XVI) are the most efficient chromosomal rearrangements for coping with high concentrations of SO_2_. In addition, we illustrate that the efficiency of the translocation VIII-t-XVI is not perfectly related to the number of 76 bp tandem repeats. Indeed, from a technological point of view the allele VIII-t-XVI^478^ would be the most efficient form to tolerate high sulfite concentrations in natural grape juice.

## Conclusion

In yeast, CR are thought to play a role in adaptation and species evolution and might have physiological consequences (Tosato and Bruschi, 2015). The setup of a *SSU1* checkup provides a rapid molecular tool for obtaining a complete overview of three CR involving the promoter region of *SSU1*. This molecular diagnostic was helpful to address some questions related to the impact of domestication of wine yeast and to progress in the identification of physio-ecological parameters that reshape the genome organization in natural isolates. From an ecological point of view, the *SSU1* checkup could be in the future a key molecular tool for addressing different questions in relation to the use of sulfites in wine. Indeed, in the recent years, the consumer-driven push for decreasing the levels of SO_2_ in the wine industry and the low-sulfite wine market are increasing and could modify the allelic frequency of *SSU1* promoter alleles. From a more applied point of view, we established a link between quantitative phenotypes and the inheritance of CR in genetically unrelated groups paving the way for yeast selection programs mediated by molecular markers.

## Acknowledgement

Authors would thank José Manuel Guillamon (Spanish National Research Council) for providing the P5 strain.

---

Figure S1. Bruvo’s distance distribution and cut off threshold used for removing very similar strains

Figure S2. Natural isolates closely related to industrial starters

Figure S3. Panel A. Principal Component analysis of 82 commercial starters discriminated by 15 polymorphic loci. The three groups represented (A to C) were inferred by k-mean clustering. The panel B represents the position of the strains according to the inferred groups.

Figure S4. Schematic representation of a VIII allele of S. cerevisiae. The figure shows the location of the 76 bp Tandem(s) and the 47 bp Tandem(s), as well as the translocation point (TP) and the upstream (5’; from the end of the forward primer to the beginning of the 76 bp Tandem) and downstream (3’; from the end of the 47 bp Tandem to the translocation point) flanking regions. The hybridization position for the primers forward (F; p1189) and reverse (R; either p1190 or p1194), are indicated.

Figure S5. Genetic distance between individuals sharing six SSU1 promoter types. The color blue and red indicate the distribution of pair wise Bruvo’s genetic distance for the total set of strains and the subset of five strains selected.

Figure S6. Growth curves of reference strains in different grape juice containing different SO2 concentrations. The data presented are the average of two independent replicates for the strain Fx10 (red), GN (green), P5 (cyan) and SB (purple). Standard error was figured out by the shaded area.

Table S1. List of the 586 strains analyzed in this work

Table S2. Additional primers used

Table S3. List of the 30 strains phenotyped

Table S4. Contingency table of natural isolates origin according to the microsatellite groups

Table S5. List of the 16 isogenic subgroups identified encompassing 125 strains

