## Supplementary material for "The *SSU1* checkup, a rapid tool for detecting chromosomal rearrangements of the *Saccharomyces cerevisiae* chromosome XVI. An ecological and technological study on wine yeast": Figure S4

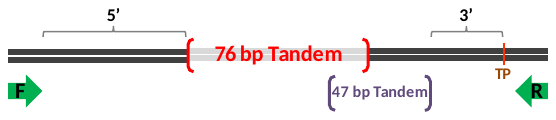

| **Alleles** | **Size (bp)^1^** | **5’ (bp)^2^** | **Tandem 76 bp^3^** | **Tandem 47 bp^4^** | **3’ (bp)^5^** |
| --- | --- | --- | --- | --- | --- |
| VIII-t-XVI | 388 | 141 | 2 | 1 | 36 |
| VIII-t-XVI | 631 | 141 | 5 | 1 | 37 |
| VIII-t-XVI | 478 | 141 | 3 | 1 | 37 |
| VIII-t-XVI | 555 | 141 | 4 | 1 | 37 |
| VIII | 615 | 141 | 1 | 1 | 48 |
| VIII | 651 | 141 | 1 | 2 | 37 |
| VIII | 649 | 141 | 1 | 2 | 37 |
| VIII | 604 | 141 | 1 | 1 | 37 |
| VIII | 667 | 141 | 2 | 1 | 37 |

^1^ PCR product size (including primers length)

^2^Size from the end of the forward primer to the beginning of the 76 bp Tandem.

^3^ Number of 76 bp Tandem in the sequence.

^4^ Number of 47 bp Tandem in the sequence.

^5^ Size from the end of the 47 bp Tandem to the translocation point.
