## Supplementary figures and images for "The *SSU1* checkup, a rapid tool for detecting chromosomal rearrangements of the *Saccharomyces cerevisiae* chromosome XVI. An ecological and technological study on wine yeast"

### Figure S1

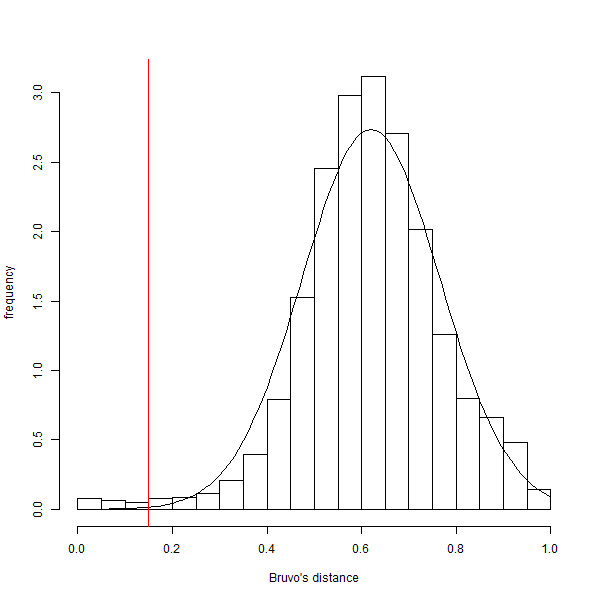

### Figure S2

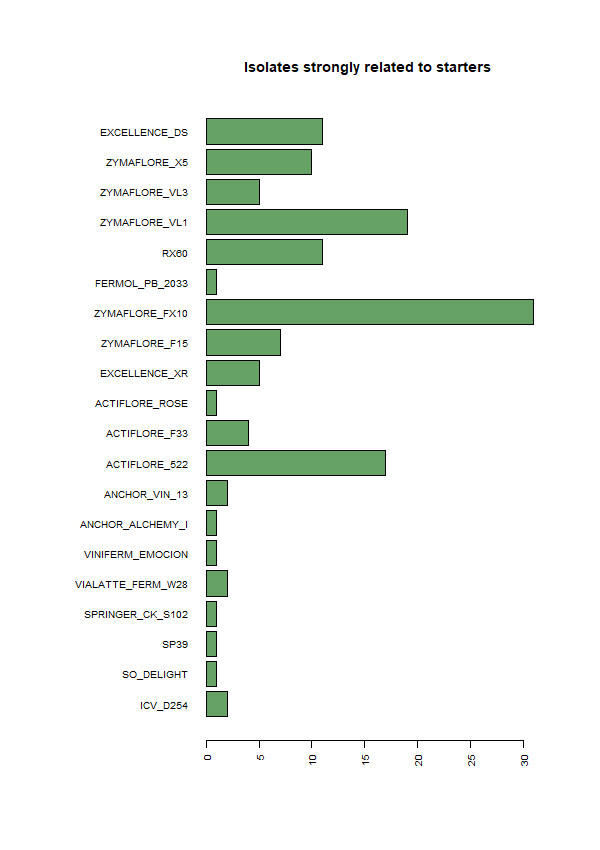

### Figure S3

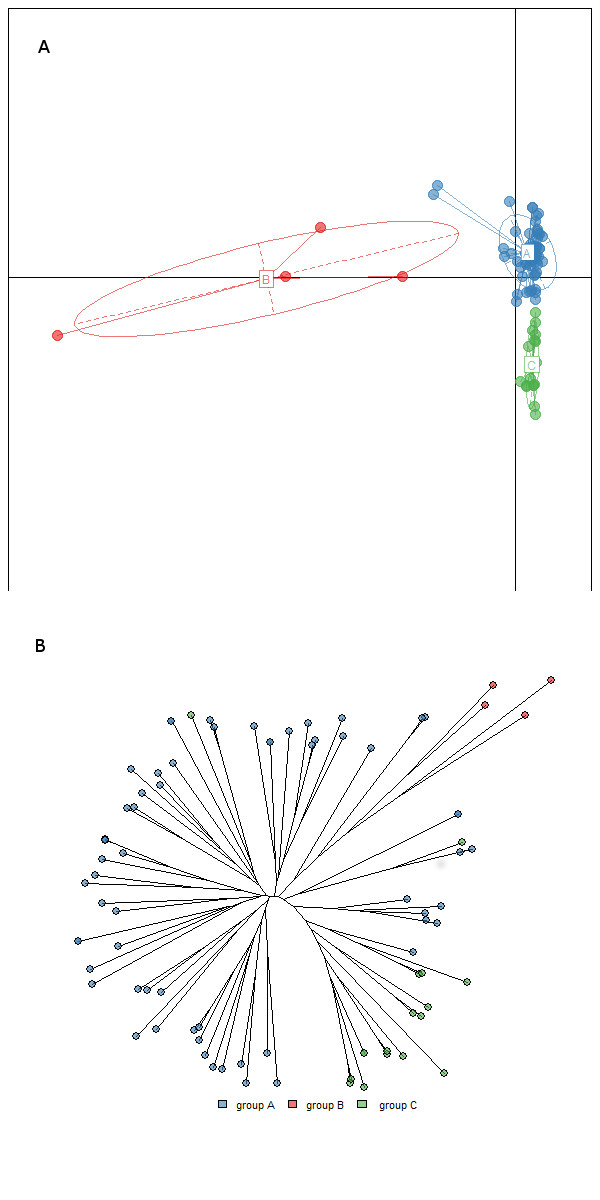

### Figure S5

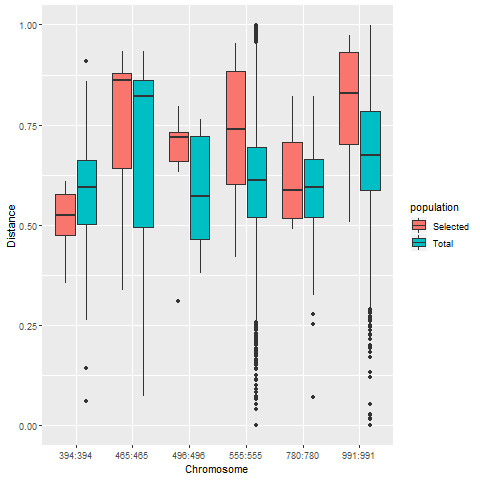

### Figure S6

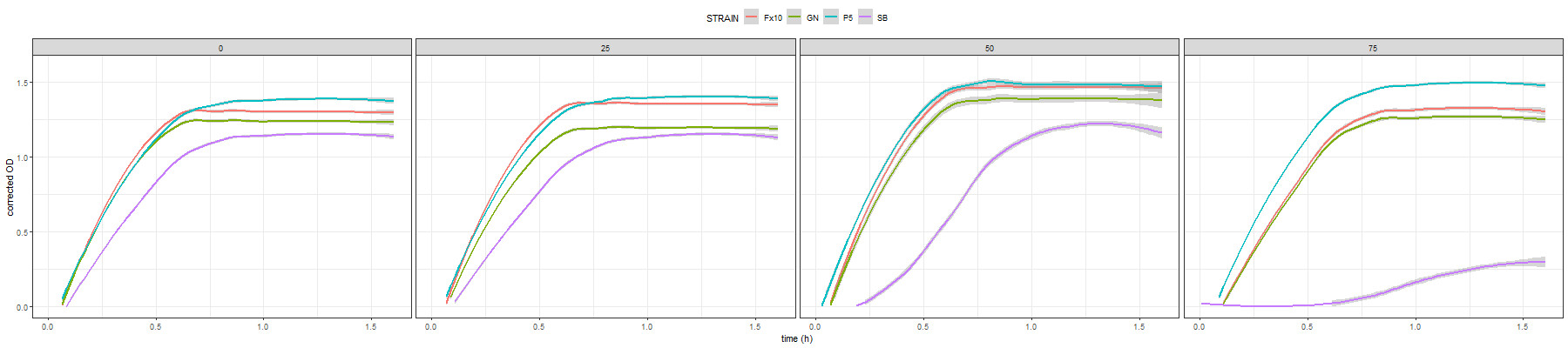
